# Sequence-level vocal convergence in common marmosets

**DOI:** 10.64898/2026.03.20.713272

**Authors:** Nakul Wewhare, Judith M. Burkart, Kaja Wierucka

## Abstract

Vocal accommodation is the process by which individuals adjust their vocalizations to resemble those of social partners. This phenomenon is widespread in social animals and can reinforce affiliation, signal group identity, and facilitate coordination. Most studies of vocal accommodation have focused on convergence in the acoustic structure of individual calls. Whether social partners also converge in how calls are arranged into sequences remains largely unknown. We examined vocal convergence during pair formation in common marmosets *(Callithrix jacchus)* by recording phee sequences from nine dyads composed of three males and three females before pairing and again four months after, in two audience contexts: when individuals interacted vocally with their partner or with an opposite sex stranger. We quantified similarity between individuals in call sequence-structure using transition probabilities, bigram frequencies, repeat-length distributions, and local alignment, and quantified similarity in acoustic structure using spectral parameters, MFCCs, and dynamic time warping. We found vocal convergence on a sequence level. After pair formation, partners became more similar in sequence structure when calling to strangers, whereas no change was detected in partner directed sequences. In contrast, call acoustic structure did not change in either context. Because vocal repertoires are constrained by anatomy and physiology, reorganizing existing call types into different combinations may provide a flexible route for modifying signals without altering the acoustic structure of individual calls. Our results provide evidence that social bonds can drive sequence level vocal convergence in a non-human primate, suggesting that vocal flexibility may arise not only through changes in acoustic structures but also through changes in how calls are organized over time.

## Introduction

Communication underpins social life by enabling animals to coordinate behaviour, negotiate conflicts, and maintain relationships across changing contexts. Among sensory modalities, vocal signals are particularly versatile: they can be produced rapidly, travel around obstacles, and be adjusted in real time to the audience and social setting (Bradbury & Vehrencamp, 2011). One manifestation of this flexibility is social vocal accommodation, where individuals adjust aspects of their vocalizations in response to affiliative partners or group members (Janik & Slater, 2000a; Ruch et al., 2018). In many species, individuals converge in the acoustic structure of their calls when social bonds form, causing partners or group members to sound more alike (Janik & Knörnschild, 2021; Ruch et al., 2018). Such convergence can reinforce social bonds, signal group identity, and help maintain cohesion (Janik & Knörnschild, 2021; Janik & Slater, 2000b; Sewall et al., 2016a).

Most evidence for social vocal accommodation in mammals concerns changes in the acoustic structure of individual calls, where individuals shift frequency contours, durations, or spectral properties so that partners sound more similar(Janik & Knörnschild, 2021; Ruch et al., 2018; Tyack, 2020). However, vocal flexibility may operate along a second, largely unexplored dimension: how calls are arranged into sequences (Kershenbaum et al., 2016). Because vocal repertoires are constrained by anatomy and physiology, reorganizing existing call types into different combinations may allow signals to change while preserving the acoustic structure of individual calls (Kershenbaum et al., 2016; Suzuki et al., 2020; Zuberbühler, 2020).

Despite growing recognition that animal vocal sequences often show non-random organization (Kershenbaum et al., 2016), whether social processes such as pair formation drive convergence at the level of call sequences remains largely untested in mammals (Kershenbaum et al., 2016; Zuberbühler, 2020). Most work examining sequence-level organization and convergence has been conducted in birds, where dialect-like variation in song syntax is well documented (Catchpole & Slater, 2008; Marler & Tamura, 1964). In contrast, evidence for socially driven reorganization of call sequences in mammals is scarce and comes mainly from studies of marine mammals (Gero et al., 2016; Hersh et al., 2022).

Common marmosets (*Callithrix jacchus*) provide an ideal system to test whether social bonds influence sequence-level vocal structure. They are socially monogamous primates that form long-term pair bonds and maintain cohesion through frequent vocal exchanges (Bezerra & Souto, 2008; Ghazanfar et al., 2026; Miller et al., 2010). Previous work has shown that newly formed pairs can converge in the acoustic properties of certain call types, suggesting that close social interaction promotes vocal similarity (Zürcher et al., 2019, 2021; Zürcher & Burkart, 2017). Marmosets produce long-distance “phee” sequences composed primarily of phee calls but often interspersed with other call types (Agamaite et al., 2015). These sequences are individually distinctive and show non-random internal organization (Bosshard et al., 2024; Gultekin et al., 2021). Yet it remains unknown whether sequence organization itself converges when social bonds form.

In many studies of vocal accommodation, animals are recorded only with their new group or partner after social bonds form, making it difficult to distinguish the potential functions of convergence (Dahlin et al., 2014; Janik & Slater, 2000b; Ruch et al., 2018). Recording individuals in different social contexts after pair formation provides an opportunity to address this question. Two main functional hypotheses have been proposed. Under the bond-reinforcement hypothesis, convergence functions as an affiliative signal and should therefore be strongest during direct interactions between partners. Under the group-signature hypothesis, convergence produces a shared vocal pattern that signals group membership or social affiliation to third parties and should therefore be most evident when interacting with non-group members (Dahlin et al., 2014; Sewall et al., 2016).

Here we tested whether pair formation induces convergence in vocal sequence structure in common marmosets and whether any convergence depends on the social audience. We compared vocal interactions of marmoset dyads before and four months after pair formation when individuals were vocalising either to their future partner or to an opposite-sex stranger. We quantified similarity at two structural levels: (1) the sequence structure of phee-call bouts and (2) the acoustic structure of individual phee calls. By comparing partner-directed and stranger-directed contexts while always measuring similarity between the same two future partners, this design allows us to test whether convergence primarily reflects bond reinforcement or the emergence of a shared pair-level vocal signature.

## Methods

### Experimental subjects

The study subjects were six captive adult common marmosets (three males and three females (Sup table 1 for more details)). Animals had continuous access to water and received a varied diet of vitamin-enriched mash and fresh fruits and vegetables twice daily, in the morning and around midday. In the afternoon, they were provided with additional protein-rich items such as insects, gum, boiled egg, or cheese. Each subject belonged to an established group or pair housed in a heated indoor enclosure (2.4 × 1.8 × 3.6 m) connected to an outdoor enclosure of (2.4 × 1.8 × 4.6 m) . All husbandry and experimental procedures complied with Swiss animal welfare regulations and were approved by the Cantonal Veterinary Office of Zürich (permit ZH232/19).

### Experimental procedure and recording

Recordings were conducted in two stages: before pair formation and after pair formation. In each stage, all nine possible male–female combinations were recorded (Figure 1a). After the first stage, three permanent dyads were established and co-housed. Four months later, the recording procedure was repeated (post-pairing stage). At each stage, six recording sessions were conducted for every male–female combination. In the post-pairing stage, recordings between bonded partners were classified as the partner context, whereas recordings between individuals that were not paired were classified as the stranger context. During a session, one male and one female were placed in adjacent wire enclosures separated by an opaque barrier that prevented visual contact while allowing acoustic communication. Each session lasted fifteen minutes. The visual barrier was present during the first and last five minutes and removed during the middle five minutes so the animals could see each other. This arrangement reliably elicited phee calling during the first and last five minutes of the session, and only phee calls produced during these periods were used for analysis. Vocalizations were recorded using two CM16/CMPA condenser microphones (Avisoft Bioacoustics), and caller identity was annotated in real time using Avisoft Recorder. For analysis, all vocal units produced by an individual within a session were treated as that individual’s session repertoire.

**Figure 1.**
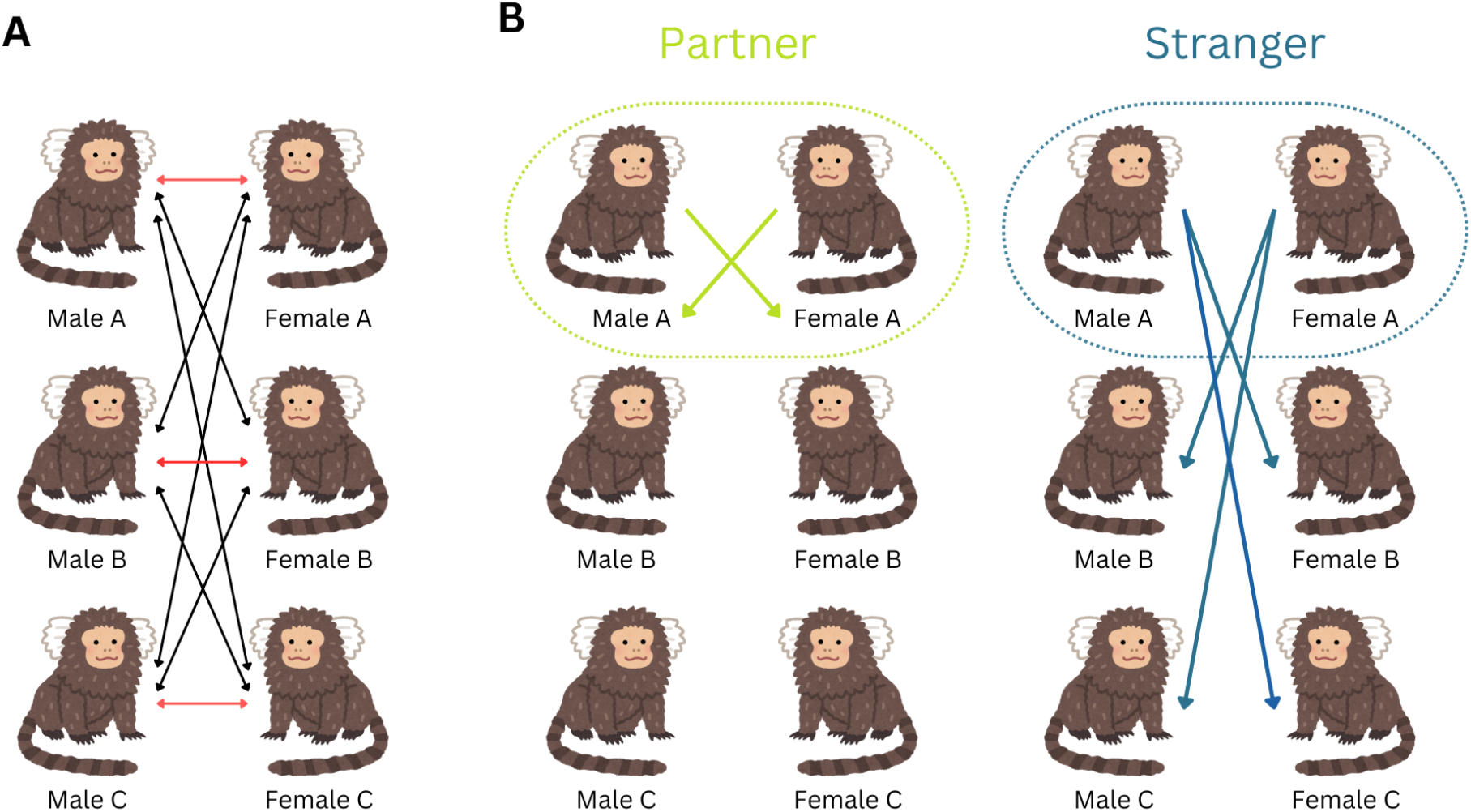
Experimental design and focal dyad comparisons. (A) Recording design across both experimental stages. In each stage, all nine possible opposite-sex male–female dyads were recorded six times. After the first stage, three permanent male–female dyads were formed and co-housed for four months before the second recording stage. Red arrows indicate the three male–female combinations that later became permanent pairs. (B) Comparison scheme for the focal dyads. Vocal similarity was always computed between the same eventual pair members. In the partner context, these two individuals were recorded while interacting with each other after pair formation. In the stranger context, those same two individuals were each recorded while interacting with opposite-sex strangers. Green outlines and arrows indicate the partner comparison; blue outlines and arrows indicate the stranger comparison.

### Data Processing

Phee calls and sequences were extracted using Avisoft-SASLab Pro version 5.3.01 (Specht, 2002). A phee sequence was defined as a bout of vocal elements separated by inter-call intervals of 0.5 s or less (Bosshard et al., 2024; Landman et al., 2020). Each sequence contained at least one phee element but could include other call types occurring within the bout. To avoid splitting a single bout at the edges, we included an isolated vocal element occurring within 1 s of the start or end of a bout when it was separated from other calls by a silence comparable to the internal inter-call intervals of that bout (Bosshard et al., 2024; Landman et al., 2020). Only calls confidently attributed to a specific individual were retained. Recordings with poor signal-to-noise ratio or overlapping calls were excluded. A 2–10 kHz bandpass filter was applied to reduce low-frequency noise. Each call was assigned to a call type by a trained annotator following Agamaite et al. (2015).

### Overview of analysis pipeline

We quantified vocal accommodation as a change in similarity between the two individuals that ultimately became pair partners. Similarity was assessed at two structural levels: sequence structure, based on the ordering of call types within phee sequences, and call structure, based on acoustic properties of individual phee calls occurring within sequences. The basic unit of analysis was a session repertoire (each pair of opposite sex individuals were recorded six times per stage), defined as all vocal units produced by one individual in one recording session. For sequence-structure analyses, the units were phee sequences. For call-structure analyses, the units were individual phee calls occurring within sequences.

Each session’s repertoire was assigned an audience context based on the identity of the conspecific across the barrier during that session. Sessions with the focal individual’s eventual partner were classified as partner sessions. Sessions with any other opposite-sex individual were classified as stranger sessions. Although all nine male–female combinations were recorded, convergence analyses focused exclusively on the three dyads that later became pair partners. Stranger-session data for a given dyad consisted of sessions in which those same individuals were recorded while in the presence of stranger individuals.

For each focal dyad, stage (before, after), and audience context (partner, stranger), we computed distances between all possible pairs of session repertoires from the two partners. Each session-pair comparison produced one distance value per metric and was retained as a separate observation for modelling. Smaller values indicate greater similarity. Within a given session-pair comparison, distances were calculated according to the metric used. For sequence alignment and spectral comparisons, this involved averaging lower-level pairwise comparisons (e.g., across sequences or calls) to produce a single distance for that session pair. These procedures are described below.

### Sequence structure analysis

We represented each vocal bout as an ordered string of discrete call-type symbols and quantified between-repertoire similarity using four complementary distance measures. Because different metrics capture different aspects of sequential organization (e.g., local motif overlap vs. global transition structure) and no single approach is uniformly optimal across animal sequence datasets, we used multiple metrics a priori and treated convergence as robust only when supported consistently across them (Kershenbaum et al., 2016; Kershenbaum & Garland, 2015).

#### 1. Transition probabilities

Animal acoustic sequences are often modelled as Markov chains (or equivalently N-gram models), where the probability of a unit depends on a finite history of preceding units; in the first-order case this is fully summarized by a transition matrix T where T_{ij} is the probability that element j follows element i. We then Vectorized the matrix, and computed Euclidean distances between vectorized transition tables, following established sequence-comparison practice. Because transition probabilities are conditioned on the preceding element, this representation emphasizes changes in conditional structure and can be sensitive to shifts involving rarer elements (i.e., it does not automatically downweight rare transitions simply because the source element is rare). (Kershenbaum & Garland, 2015).

#### 2. Bigram distribution

We computed the normalized frequency distribution of all two-element subsequences (bigrams) and compared repertoires via Euclidean distance between bigram proportion vectors. N-gram models are widely used for animal sequences and provide a complementary summary of sequential patterning; unlike transition probabilities, bigrams normalize by total bigram count rather than by the frequency of the source element, thereby reflecting overall usage of two-element chunks. Kershenbaum & Garland, 2015).

#### 3. Repeat distribution of phee

Motivated by the prevalence of repetition in our data, we computed the count for the number of repetitions for phee elements and used the Euclidean distance between repeat probability distributions.

#### 4. Local alignment

Because our bouts vary in length, comparing whole sequences end-to-end can be inappropriate; instead, identifying and scoring the best-matching subsections can better capture shared motifs embedded within longer bouts ( Dhati et al., 2025; Kershenbaum & Garland, 2015). We therefore used Smith–Waterman local alignment, which finds the highest-scoring locally matched region rather than forcing a global alignment of entire sequences (Smith & Waterman, 1981). We used match/mismatch/gap scores of 1/-2/-2 to favor strict local matches, and applied a length-based normalization (dividing by the log of matched-sequence length) to reduce spurious high scores when short sequences are matched against very long sequences (Dhati et al., 2025). To compare repertoires, we computed best-match scores for each focal sequence against all sequences in the other repertoire, averaged these best scores, repeated with focal/matched roles reversed to ensure symmetry, and converted similarity to a distance by multiplying by -1 (Dhati et al., 2025).

### Call structure analysis

Call-structure analyses were performed on phee calls occurring within sequences.

#### 1. Spectro-temporal parameters (STP)

Energy-based spectral and duration parameters were extracted in *Raven Pro 1*.*6*.*5* using spectrograms generated with a Hann window (400 ms), 200 ms hop size, and 50% overlap. Extracted parameters included frequency percentiles, center frequency, bandwidth measures, duration 50%, and entropy measures (Odom et al., 2021). After z-normalization, principal component analysis was performed, and the first five components explaining approximately 97% of variance were retained. Euclidean distances were computed in this reduced space.

#### 2. Mel-frequency cepstral coefficients (MFCC)

To calculate MFCCs, we used the package ‘tuneR’ (Ligges et al., 2004) in the R programming environment (R Core Team, 2025). We extracted 133 MFCC (Clink & Klinck, 2021) from the same selections and performed a separate PCA after z-normalization, retaining the first five principal components, which explained about 76% of variance. Euclidean distances between calls in this space provided a second spectral distance. MFCC’s have been shown to be better than historically used STP, and have been shown to be great at deciphering fine scale differences across taxa (Wierucka et al., 2025).

#### 3. Dynamic time warping (DTW)

We implemented a spectrogram based DTW that compares columns using cosine distance and finds an optimal alignment path (Sakoe & Chiba, 1978). To remove bias from call duration, we normalized the DTW cost by the sum of the lengths of the two spectrograms. This produces a distance that reflects similarity in frequency modulation patterns rather than length, and has been extensively used in analysis of human speech (Müller, 2007). As shown by recent studies we do believe that differences in the phee calls across individuals are among the frequency modulation patterns used (Oren et al., 2024), thus we used DTW which has been shown to be especially good at capturing differences in frequency modulation while controlling for slight differences in duration (Müller, 2007).

For each spectral metric, dyadic repertoire distance was calculated as the mean of all pairwise distances between phee calls from the two partners’ session repertoires. These distances between the two partners’ session repertoires were used in our models.

### Modelling vocal distance change

All analyses were conducted in R (R Core Team, 2025) using Bayesian multilevel models implemented using R package *brms* (Bürkner, 2018). For each dataset (sequence–partner, sequence–stranger, call–partner, call–stranger), we modelled partners’ session repertoire distances as a function of the experimental stage (before versus after pairing). The distances were z normalized within a distance metric, but across pairs, stage (before, after), and audience context (partner, stranger)

Distances were continuous and approximately symmetric after z-scoring; therefore, we used Gaussian error models. Because distance values were derived from multiple pairwise comparisons between session repertoires of the same two individuals, observations were not independent (Hart et al., 2022). Each dyad contributed multiple session-pair combinations per stage and context. To account for this hierarchical structure, we included random intercepts for pair identity, focal session, and partner session (Bürkner, 2018; Hart et al., 2022). Including focal and partner sessions as random effects accounts for repeated use of the same session repertoires across multiple pairwise distance calculations.

For pooled models integrating multiple distance metrics within a structural level, we included metric as an additional random intercept. This treats metrics as repeated measures of the same underlying construct (sequence or call distance) while allowing baseline differences across metrics. To ensure that pooled conclusions were not driven by any single metric, we additionally fitted separate per-metric models as robustness checks.

Stage was coded as a centred dummy variable (before = −0.5, after = +0.5). Centred coding improves sampling efficiency in multilevel models and ensures that the intercept corresponds to the grand mean across stages (Gelman & Hill, 2007; McElreath, 2020). Under this coding scheme, the stage coefficient β represents the standardized after–before change in distance. Because distances were z-scored within the metric, β is expressed in standard deviation units. A negative β indicates a reduction in distance after pairing (convergence), whereas a positive β indicates an increase in distance (divergence).

Models were fitted with four MCMC chains, each with 2,000 warm-up and 2,000 sampling iterations, using weakly regularising priors (Normal(0,1) for fixed effects; Exponential(1) for standard deviations) (Gelman et al., 2013). Model adequacy was assessed using posterior predictive checks (McElreath, 2020), convergence diagnostics. Across pooled models, R-hat values ranged from 1.000 to 1.004, effective sample sizes were high (minimum bulk ESS = 1273; minimum tail ESS = 1663), no divergent transitions were observed, and Pareto-k values were ≤ 0.694 indicating the model results should be reliable (Gelman et al., 2013; McElreath, 2020). Posterior predictive plot was used to make sure the model can replicate data distribution faithfully in its predictions. Inference is reported using posterior medians and 95% credible intervals for the stage effect (Gelman et al., 2013; McElreath, 2020).

## Results

We analysed 3,904 individual calls for spectral comparisons and 1,619 phee sequences for sequence analyses. Most sequences were composed primarily of phee elements (72%), although other call types such as peeps, eggs, tsks, ocks, trillphees, and sd-peeps also occurred (Figure 2A). Sequences were generally short, with a mean of 3.21 (± 1.5 SD) calls per sequence, and 82.7% contained four or fewer elements (Figure 2B). Self-transitions were common, indicating a strong tendency toward repetition (Figure 2C). Certain call types co-occurred more frequently than others (Figures 2C and 2D), consistent with structured, non-random sequence organization reported in previous studies.

**Figure 2.**
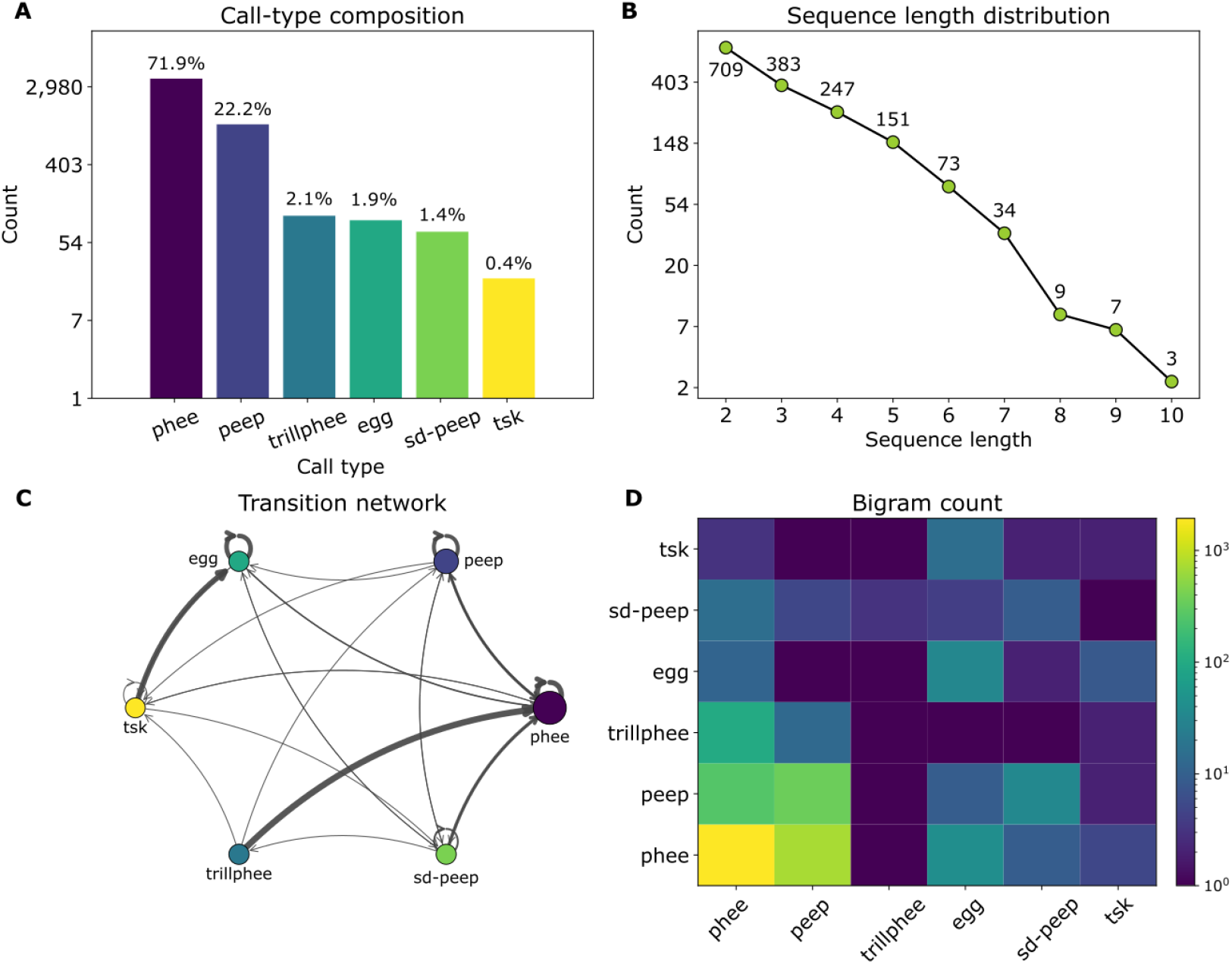
Structural features of common marmoset phee sequences. Summary of 1,619 phee sequences analysed across all individuals. (A) Relative abundance of call types within sequence. The y-axis is shown on a natural-log scale. (B) Distribution of sequence lengths, where counts are plotted on a natural-log scale. (C) Transition network, where nodes represent the call type and arrows represent the transition probability between them. Arrow thickness is proportional to the transition probability between call types. (D) Bigram count matrix showing the relative frequency of all two-element combinations (columns = first element, rows = second element).

### Sequence structure

Pair formation was associated with a decrease in sequence-structure distance when individuals called to strangers. After pairing, sequence repertoires were more similar in the stranger context (β = −0.983, 95% CrI [−1.375, −0.604], p = 0.000, N = 500; Figure 3). In contrast, no change in sequence-structure distance was detected in the partner-directed context (β = 0.235, 95% CrI [−0.545, 0.986], p = 0.526, N = 248; Figure 3).

**Figure 3.**
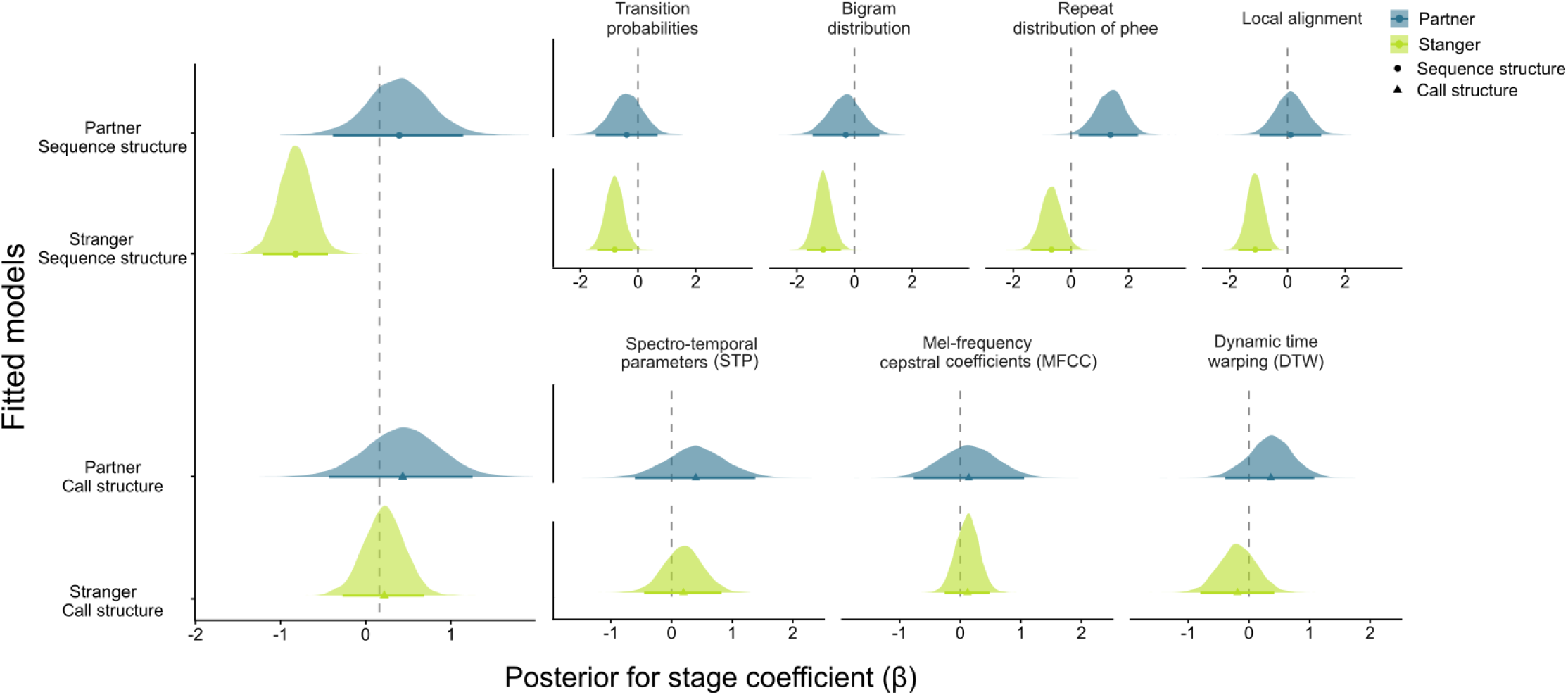
Context- and structure-specific vocal accommodation in common marmosets. Posterior distributions of the stage effect (β; after − before) from Bayesian multilevel models estimating changes in repertoire distance following pair formation. Results are shown separately for sequence structure (top panels) and call structure (bottom panels), and for partner-directed (blue) and stranger-directed (green) contexts. The leftmost panels show pooled models across metrics within each structural level; remaining panels show metric-specific models. Negative β values indicate convergence (reduced distance after pairing), whereas positive values indicate divergence. Points denote posterior medians and shaded areas represent 95% credible intervals. The dashed vertical line marks zero, corresponding to no change across stages.

Per-metric models broadly supported this pattern. In the stranger context, all four sequence metrics detected reductions in distance after pairing: transition probabilities (β = −0.812, 95% CrI [−1.417, −0.186], p = 0.010, N = 125), bigram distribution (β = −1.084, 95% CrI [−1.667, −0.469], p = 0.001, N = 125), and local alignment (β = −1.123, 95% CrI [−1.708, −0.552], p = 0.000, N = 125). The repeat distribution (β = −0.680, 95% CrI [−1.393, 0.041], p = 0.064, N = 125; Figure 3). In the partner context, no consistent reduction in distance was detected across metrics; the repeat distribution showed an increase in distance after pairing (β = 1.369, 95% CrI [0.270, 2.325], p = 0.018, N = 62), while transition probabilities (β = −0.392, 95% CrI [−1.465, 0.676], p = 0.468, N = 62), bigram distribution (β = −0.305, 95% CrI [−1.452, 0.859], p = 0.581, N = 62), and local alignment (β = 0.108, 95% CrI [−0.964, 1.172], p = 0.833, N = 62) showed no clear change (Figure 3).

### Call structure

Pair formation was not associated with a reduction in call-structure distance in either context. In the stranger condition, call-structure distance did not change after pairing (β = 0.059, 95% CrI [−0.433, 0.524], p = 0.809, N = 429; Figure 3). In the partner condition, no change was detected (β = 0.276, 95% CrI [−0.593, 1.095], p = 0.520, N = 228; Figure 3).

Per-metric spectral models were consistent with this pattern. In the stranger context, none of the spectral metrics detected a change after pairing (STP: β = 0.195, 95% CrI [−0.453, 0.826], p = 0.540, N = 143; MFCC: β = 0.121, 95% CrI [−0.260, 0.488], p = 0.544, N = 143; DTW: β = −0.188, 95% CrI [−0.800, 0.420], p = 0.552, N = 143). In the partner context, none of the spectral metrics detected a change after pairing (STP: β = 0.398, 95% CrI [−0.605, 1.381], p = 0.413, N = 76; MFCC: β = 0.140, 95% CrI [−0.767, 1.054], p = 0.760, N = 76; DTW: β = 0.364, 95% CrI [−0.392, 1.076], p = 0.328, N = 76; Figure 3).

## Discussion

Our research showed that pair formation in common marmosets was associated with convergence in the sequence structure of phee sequences, suggesting that partners develop a shared combinatorial pattern in how calls are arranged over time. Interestingly, we found this sequence-level convergence only when individuals vocalised to strangers, which is consistent with the group signature hypothesis, whereby individuals signal their social affiliation through shared vocal patterns expressed toward third parties (Dahlin et al., 2014; Fischer & Hammerschmidt, 2020; Ruch et al., 2018; Sewall et al., 2016b). In contrast, we did not detect systematic changes in the acoustic structure of individual phee calls across contexts. Given that phee calls function as individually distinctive long distance contact calls, convergence in sequence organization may provide a way for partners to signal their social bond while maintaining stable acoustic cues to individual identity (Ghazanfar et al., 2026; Oren et al., 2024; Zürcher et al., 2021). To our knowledge, this is the first evidence of sequence-level accommodation in a nonhuman primate, highlighting that sequence combinatorial organization is not only functional but can also be shaped by social relationships.

The fact that convergence occurred only in the stranger context supports the interpretation that partners develop a shared sequence structure that functions as a pair level signal when interacting with others. Under this interpretation, sequence convergence allows individuals to advertise their social affiliation without compromising the individual identity information encoded in acoustic structures. A similar division between stable acoustic elements and socially structured temporal patterns occurs in other taxa. In sperm whales, for example, social identity is encoded primarily in the temporal structure of click sequences rather than in the acoustic properties of individual clicks (Gero et al., 2016; Hersh et al., 2022; Rendell & Whitehead, 2003). Likewise, socially guided convergence in sequence patterns has been documented in budgerigars when previously separated groups interact (Madabhushi et al., 2023). Across these systems, reorganization of stable vocal units appears to provide a flexible mechanism for signalling social relationships while preserving individual acoustic signatures.

Our findings add to growing evidence that primate vocal systems make flexible and versatile use of call combinations rather than relying solely on isolated cues. In marmosets, call sequences show non-random organization with higher-order dependencies beyond simple pairwise transitions (Bosshard et al., 2024), and individuals differ systematically in how they structure these sequences across development (Gultekin et al., 2021). Our results extend this work by showing that sequence structure is not only individually patterned but can also shift with changes in social relationships. Studies in chimpanzees demonstrate that individuals produce diverse vocal sequences in which call types are flexibly recombined into ordered structures (Girard-Buttoz, Bortolato, et al., 2022; Girard-Buttoz, Zaccarella, et al., 2022), and that such combinations can expand communicative meaning through multiple combinatorial mechanisms (Girard-Buttoz et al., 2025). In bonobos, some call combinations even show non trivial compositional structure in which the meaning of one element modifies the meaning of the other element in the sequence (Berthet et al., 2025). Because primate vocal repertoires are constrained by anatomy and physiology, recombining a limited set of calls into different sequences may provide an efficient route for increasing signal diversity without requiring the evolution of new call types (Kershenbaum et al., 2016; Suzuki et al., 2020; Zuberbühler, 2020). In this light, the sequence-level convergence we observe in marmosets indicates that this combinatorial organization is not only structured and meaningful but can also be shaped by social relationships.

The absence of call level convergence is equally informative. Phee calls are long distance contact calls known to encode caller identity and sex (Meshinska et al., 2024; Miller et al., 2010; Oren et al., 2024; Phaniraj et al., 2023). Strong convergence in acoustic structure could therefore risk eroding identity cues that are important for reliable recognition over distance. Previous work has similarly reported weaker or inconsistent convergence in phee calls compared with other marmoset call types (Snowdon & Elowson, 1999; Zürcher et al., 2021). An additional explanation may be baseline similarity. Because all individuals in our study originated from the same colony, they may already have shared similar acoustic variants, leaving less scope for further spectral convergence (Zürcher et al., 2019, 2021). These findings suggest that acoustic structure and sequence organization may be shaped by different functional constraints.

Although the number of dyads was modest, the longitudinal within-dyad design and the consistency of the effect across multiple sequence metrics provide convergent support for sequence-level convergence after pair formation within the present sample. Replication with a larger number of individuals will nevertheless be important for determining how broadly this pattern generalises. While we demonstrated that convergence emerged specifically in the stranger context, suggesting that these sequence-level changes are socially relevant, these conclusions are based on production data alone, and future studies should incorporate playback or interactive experiments to test directly whether marmosets perceive and use these cues during social interactions.

Our results identify sequence organisation as an important and previously underappreciated dimension of primate vocal accommodation. Work on vocal flexibility has focused largely on changes in the acoustic structure of individual calls, however our findings show that social relationships can also reshape how stable call types are arranged over time. In species whose vocal repertoires are constrained by anatomy and physiology, such combinatorial reorganisation may provide an efficient route for generating socially meaningful variation without sacrificing individual recognisability. By linking pair formation to changes in call ordering, this study helps connect vocal accommodation with emerging work on mammal call combinations and underscores the importance of analysing sequence structure alongside the acoustic structure of vocalisations.

## Acknowledgement

We thank Monika Mircheva, Adele Tuozzi and Kristin Meshinska for data collection. We thank Elliot Howard-Spink and Nikhil Phaniraj for thoughtful feedback on the manuscript.

## Data Availability and code

R and Python scripts along with supporting data can be found at https://github.com/Nakul-wewhare/Marmoset-phee-sequence-accodomation

## Supplementary

**Table S1.** Subjects included in the study. The table lists the six common marmosets (Callithrix jacchus) used in the experiment, including their sex, subject ID, and date of birth. Additional information indicates known kin relationships or prior housing conditions relevant to the experimental setup.

| Sex | Subject ID | Date of Birth | Additional Information |
| --- | --- | --- | --- |
| Male | A | 30.10.2008 | Following the death of his previous partner, he was housed with his son (subject B) before the experiment |
| Male | B | 23.10.2019 | Son of subject A |
| Male | C | 25.02.2016 | Brother of subject F |
| Female | D | 29.03.2013 | – |
| Female | E | 18.11.2019 | – |
| Female | F | 10.01.2018 | Sister of subject C |

